# *Haemophilus* to meningococci transfer of beta-lactamase

**DOI:** 10.1101/308056

**Authors:** Eva Hong, Ala-Eddine Deghmane, Muhamed-Kheir Taha

**Affiliations:** Invasive Bacterial Infection and National Reference Centre for meningococci, Institut Pasteur, Paris, France

## Abstract

We report the detection in France of a beta-lactamase producing invasive meningococcal isolate. Whole genome sequencing of the isolate revealed ROB-1 type beta-lactamase that is frequently encountered in *Haemophilus influenzae* suggesting horizontal transfer between isolates of these bacterial species. Beta-lactamases are exceptional in meningococci with no reports from more than two decades. This report is worrying as the expansion of such isolates may jeopardize the effective treatment against invasive meningococcal disease.

---

Beta lactam antibiotics should be administered upon suspicion of invasive meningococcal disease (IMD). Reduced susceptibility to penicillin G, ampicillin and amoxicillin has been increasingly reported worldwide. Isolates with reduced susceptibility to third generation cephalosporins have been also described through gene transfer from gonococci to meningococci. However, beta-lactamase-producing meningococci remained exceptional with no reported cases for more than two decades (1).

We identified in 2017 at the national reference centre for meningococci in Paris, France, an invasive isolate of *N. meningitidis* that produced a beta-lactmase. The isolate was from the blood of a 23-year old woman. Antibiotic susceptibility testing revealed resistance to penicillin G and amoxicillin with minimal inhibitory concentrations (MIC) of 3 and 12 mg/L respectively. The clavulanic acid (an inhibitor to beta-lactamase) decreased the amoxicillin MIC to 0.5 mg/L. The isolated was susceptible to third generation cephalosporins (ceftriaxone and cefotaxime, MIC=0.008 mg/L) and the patient was successfully treated with ceftriaxone. The isolate was of serogroup Y. Whole genome sequencing (WGS) allowed genotyping of the isolate [group: PorA variable region VR1, PorA variable region VR2: FetA variable region: clonal complex (Sequence type)]. The isolate showed the combination Y:P1.5-2, 10-2:F4-1:cc23 (ST-3587). The data (Fasta Format) are accessible on the PubMLST web site (accession number (ID) 53820, isolate, LNP29202) and through the NCBI GenBank accession N° PRJNA454456. WGS also allowed detection of a ROB-1 type beta lactamase (*bla*_ROB-1_) on the chromosome of the isolate and that was 100% identical at both DNA and amino-acid levels to *bla*_ROB-1_ harbored by the pB1000 in *Haemophilus influenzae* but also found in other bacterial species such as *Pasteurella* and *Moraxella* species (2). The sequence analysis showed that the *bla*_ROB-1_ gene was inserted on the chromosome of the LNP29202 isolate upstream the gene Neis0803 and downstream the gene Neis0807. These two genes encode hypothetical proteins. The *bla*_ROB-1_ gene was present on the meningococcal chromosome with its promoter as well as the ribosome binding site allowing transcription and translation of the beta lactamase. The *bla*_ROB-1_ gene of the plasmid pB1000 was previously shown to be functional and responsible for the resistance to beta lactamas antibiotics (3). However, no additional genes from the pB1000 (such as the *mobABC* genes) were transferred with the *bla*_ROB-1_ gene on the chromosome of the LNP29202 isolate (Fig. 1). It has been previously suggested that the flanked sequences of the *bla*_ROB-1_ gene on the pB1000 plasmid mediates the insertion of this gene. Indeed, a perfect two repeats of the sequence GACTT that are linked to a transcriptional terminator, are located upstream and downstream the *bla*_ROB-1_ on the pB1000 plasmid (3). The *bla*_ROB-1_ in the LNP29202 isolate is preceded by only one copy of the GACTT that was linked to a transcriptional terminator as in the plasmid pB1000 (Fig.1). A similar organization was also reported in the plasmid pHS-Tet encoding tetracycline resistance in *Haemophilus parasuis* strain HS1543 (4). The acquisition of *bla*_ROB-1_ gene may have occurred through a respiratory co-infection of *N. meningitidis* and isolates of these bacterial species. The isolates of the clonal complex cc23 are increasing in France and in other European countries since 2010(5). They are frequently involved in pneumonia particularly in elderly (6). However, the median age of the IMD due to cc23 isolates seems to shift to younger ages. It was significantly (p=0.0003) lower than that for the other cc in France for the period 2012-2017 (Figure). This shift to younger ages, where meningococcal carriage is frequent(7), increases the frequency of horizontal gene transfer between meningococcal isolates and isolates of other bacterial species allowing the acquisition and the spread of resistance mechanisms. The acquisition of ROB-1 beta-lactamase by serogroup Y-cc23 isolates is therefore worrying due to the shift to younger ages where the carriage and transmission occur at high rate (7). Preventive strategies (vaccination) should be enhanced to reduce the incidence of the disease and the circulation of the isolates. These measures should reduce the use of antibiotics and selection of resistance.

**Figure 1.**
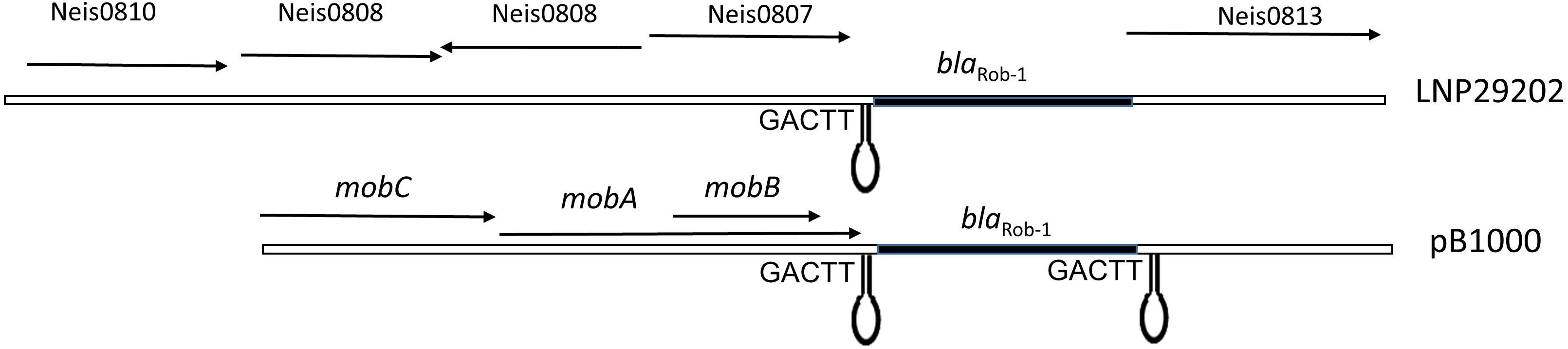
Schematic representation of the genetic organization of the chromosomal region of the isolate LNP29202 that received the insertion of the *bal_ROB-1_* gene. The organization of the pB1000 plasmid harboring the *bal_ROB-1_* gene is also shown. The arrow indicated the genes (names are above the arrows). The GACTT sequences and the transcription terminator (the hairpin structure) are also indicated.

**Figure 2.**
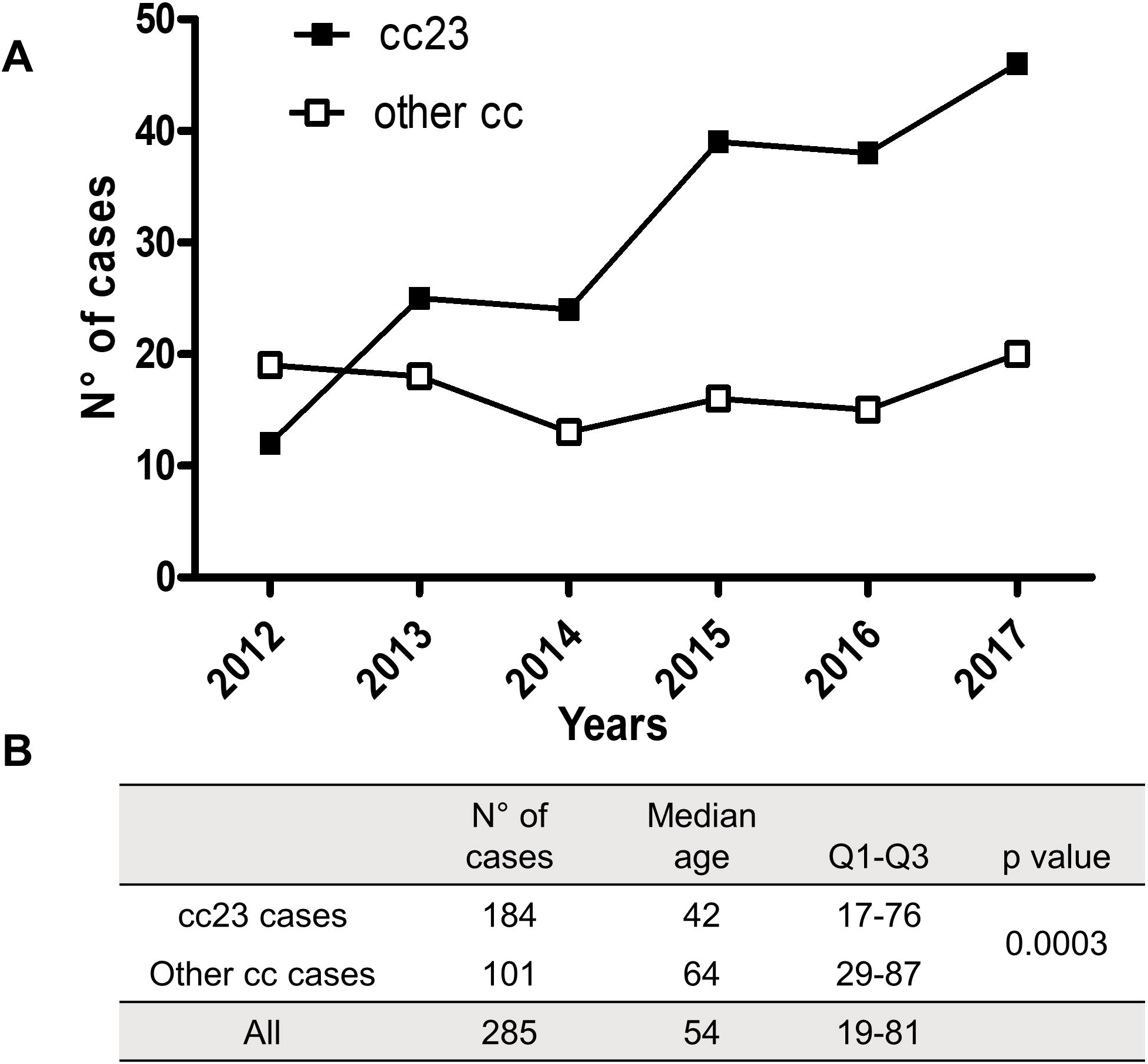
(A) Evolution of cases (numbers) of group Y invasive meningococcal disease in France per year during the period 2012-2017 according to clonal complexes (cc23 versus other cc). (B) Evolution of cases (numbers) of IMD for the whole period according to median age and to the clonal complexes (cc23 versus other cc). A chi-squared test was performed to calculate p value (cc23 versus other cc).

## Acknowledgements

This work made use of the Neisseria Multi Locus Sequence Typing website (https://pubmlst.org/neisseria/) developed by Keith Jolley and sited at the University of Oxford (8). The development of this site has been funded by the Wellcome Trust and European Union. We also acknowledge the PIBNET-P2M platform at the Institut Pasteur.

## References

1. Deghmane AE, Hong E, Taha MK. 2017. Emergence of meningococci with reduced susceptibility to third-generation cephalosporins. J Antimicrob Chemother 72:95–98.

2. Tristram SG, Littlejohn R, Bradbury RS. 2010. blaROB-1 presence on pB1000 in Haemophilus influenzae is widespread, and variable cefaclor resistance is associated with altered penicillin-binding proteins. Antimicrob Agents Chemother 54:4945–7.

3. San Millan A, Escudero JA, Catalan A, Nieto S, Farelo F, Gibert M, Moreno MA, Dominguez L, Gonzalez-Zorn B. 2007. Beta-lactam resistance in Haemophilus parasuis Is mediated by plasmid pB1000 bearing blaROB-1. Antimicrob Agents Chemother 51:2260–4.

4. Lancashire JF, Terry TD, Blackall PJ, Jennings MP. 2005. Plasmid-encoded Tet B tetracycline resistance in Haemophilus parasuis. Antimicrob Agents Chemother 49:1927–31.

5. Broker M, Emonet S, Fazio C, Jacobsson S, Koliou M, Kuusi M, Pace D, Paragi M, Pysik A, Simoes MJ, Skoczynska A, Stefanelli P, Toropainen M, Taha MK, Tzanakaki G. 2015. Meningococcal serogroup Y disease in Europe continuation of high importance in some European regions in 2013. Hum Vaccin Immunother:0.

6. Sall O, Stenmark B, Glimaker M, Jacobsson S, Molling P, Olcen P, Fredlund H. 2017. Clinical presentation of invasive disease caused by Neisseria meningitidis serogroup Y in Sweden, 1995 to 2012. Epidemiol Infect 145:2137–2143.

7. Christensen H, May M, Bowen L, Hickman M, Trotter CL. 2010. Meningococcal carriage by age: a systematic review and meta-analysis. Lancet Infect Dis 10:853–61.

8. Jolley KA, Maiden MC. 2010. BIGSdb: Scalable analysis of bacterial genome variation at the population level. BMC Bioinformatics 11:595.

